# The Role of TandemHeart^TM^ combined with ProtekDuo^TM^ as Right Ventricular Support Device: A Simulation Approach

**DOI:** 10.1101/2024.07.29.604074

**Authors:** Beatrice De Lazzari, Roberto Badagliacca, Massimo Capoccia, Marc O Maybauer, Claudio De Lazzari

## Abstract

Right ventricular failure increases short-term mortality in the setting of acute myocardial infarction, cardiogenic shock, advanced left-sided heart failure and pulmonary hypertension. Right ventricular failure remains quite a challenging condition to manage in view of its complex background and still incomplete understanding of its pathophysiology. Percutaneous and surgically implanted right ventricular assist devices (RVADs) have been investigated in different clinical settings. The use of the ProtekDuo^TM^ (LivaNova, London, UK) is currently a promising approach due to its features such as groin-free approach leading to early mobilisation, easy percutaneous deployment, compatibility with different pumps and oxygenators, and adaptability to different configurations. The aim of this work was to simulate the behaviour of the TandemHeart^TM^ pump applied “*in series*” and “*in parallel*“ mode and the combination of TandemHeart^TM^ and ProtekDuo^TM^ cannula as right ventricular assist device using CARDIOSIM^©^ software simulator platform. The effects induced on the main hemodynamic and energetic variables were analysed for both the right atrial-pulmonary arterial and right ventricular-pulmonary arterial configuration with different pump rotational speed and following of Milrinone administration. The TandemHeart^TM^ increased right ventricular end systolic volume by 10%, larger increases were evident for higher speeds (6000 and 7500 rpm) and connections with 21 Fr inflow and 17 Fr outflow cannula, respectively. Both TandemHeart^TM^ and ProtekDuo^TM^ support increased left ventricular preload. When different RVAD settings were used, Milrinone therapy increased the left ventricular pressure-volume area and decreased the right pressure-volume area slightly. A reduction in oxygen consumption (demand) was observed with reduced right stroke work and pressure volume area and increased oxygen supply (coronary blood flow).

## Introduction

Right ventricular failure increases short-term mortality in the setting of acute myocardial infarction, cardiogenic shock, advanced left-sided heart failure and pulmonary hypertension. Underlying mechanisms consist of contractile failure following myocardial ischaemia or inflammation caused by myocarditis; volume overload due to right-sided valvular regurgitation; volume overload secondary to increased venous return or displacement of the interventricular septum towards the left ventricle (LV) after insertion of a left ventricular assist device (LVAD); pressure overload related to decompensated left-sided heart failure, worsening pulmonary hypertension and acute pulmonary embolism [1]. Right ventricular failure remains quite a challenging condition to manage in view of its complex background and still incomplete understanding of its pathophysiology [2-4]. Medical treatment optimisation in the acute setting includes targeted volume administration, vasopressors to support coronary perfusion, inotropes to restore contractility and pulmonary vasodilators to reduce afterload. Percutaneous and surgically implanted right ventricular assist devices (RVADs) have been investigated in different clinical settings [5]. The use of the ProtekDuo^TM^ (LivaNova, London, UK) is currently a promising approach in this context due to its features [6]. The device is a single-site, dual-lumen cannula for percutaneous insertion through the right internal jugular vein and advanced to sit in the main pulmonary artery (PA), approximately 1-2 cm distally from the pulmonary valve, under echocardiographic [7,8] or fluoroscopic guidance [9]. The proximal fenestration of the cannula drains venous blood from the right atrium to the extracorporeal membrane oxygenation (ECMO) system and re-infuses it in the main pulmonary artery through the distal fenestration. This configuration, namely V-P ECMO, bypasses the right ventricle making the ProtekDuo^TM^ a suitable and effective right ventricular assist device [10] resulting in improved pulmonary flow, left atrial filling pressure and left ventricular preload. Consequently, ventricular interdependence will be addressed by reducing the leftward shift of the interventricular septum observed in right ventricular failure [11]. Originally designed as right ventricular assist device (RVAD) cannula to be used with the TandemHeart^TM^ (LivaNova PLC, London, UK) [12-14] and subsequently with a centrifugal pump [16], the ProtekDuo^TM^ has been used in different ECMO configurations in the context of Covid-related ARDS, right ventricular and bi-ventricular failure [16-19]. The use of the ProtekDuo^TM^ as RVAD has several advantages compared to other RVAD systems [6,18] and allows a flexible approach due to its versatility [20-22].

The ProtekDuo^TM^ device has significant potential in view of its versatility of use such as groin-free approach leading to early mobilisation, easy percutaneous deployment, compatibility with different pumps and oxygenators, and adaptability to different configurations [21] and easy weaning with bedside decannulation [23].

The aim of this work was to reproduce the behaviour of the TandemHeart^TM^ pump and the combination of TandemHeart^TM^ and ProtekDuo^TM^ cannula as right ventricular assist device (RVAD) using CARDIOSIM^©^ software simulator platform [24]. To achieve this, two new modules have been implemented in the software. The first module simulated the TandemHeart^TM^ pump in RVAD configuration (TH-RVAD), both as a right atrial-pulmonary arterial and a right ventricular-pulmonary arterial connection, driven by four different rotational speeds. The second module reproduced the behaviour of the ProtekDuo^TM^ cannula plus TandemHeart^TM^ (TH-PD-RVAD). Placement of the outflow of the ProtekDuo^TM^ in the main pulmonary artery (PA) allows a complete bypass of the failing right ventricle improving pulmonary flow, left atrial filling pressures (LAP), and left ventricular (LV) preload. This can also reduce ventricular interdependence and leftward shift of the interventricular septum that occurs in right ventricular failure (RVF).

The effects induced by the TandemHeart^TM^ on the main hemodynamic and energetic variables were analysed for both the right atrial-pulmonary arterial (RA-PA) and right ventricular-pulmonary arterial (RV-PA) configuration with different pump rotational speed and with different size of connecting cannulae. In addition, the effects induced by the TH-PD-RVAD on the same variables were analysed and discussed.

## Material and Methods

CARDIOSIM^©^ is the software simulator platform of the cardiovascular system, mechanical circulatory and ventilatory supports developed in the Cardiovascular Numerical/Hybrid Modelling Lab of the CNR Institute of Clinical Physiology (Rome). The modular logic of CARDIOSIM^©^’s design enables the combination of various modules to create various cardiovascular network configurations for the haemodynamic analysis of specific systems based on numerical models of variable complexity. For example, coupling of ventricular and arterial elastance with changes in the parameters of the left ventricle and/or the systemic arterial branches can be studied by simulating the behaviour of the left network only, which consists of atrium, ventricle and systemic circulation [25].

The present work is based on the implementation of two new modules within CARDIOSIM^©^ software platform: the first replicates the behaviour of the TandemHeart^TM^ pump whilst the second replicates the behaviour of the ProtekDuo^TM^ cannula. These new settings will allow a detailed investigation of the devices interactions and their influence on the haemodynamic and energetic variables of the cardiovascular system.

The TandemHeart^TM^ pump module consists of an inflow (drainage) from the right atrium (RA) and an outflow (return) into the main pulmonary artery (PA) according to [26]. Equation (1) simulates the experimental “pressure drop-flow rate” waveforms for the right atrial/ventricular drainage cannula (21 Fr) and the arterial (outflow) cannula (12 Fr and/or 17 Fr). The behaviour of the pump when running at four different rotational speeds (3000, 4500, 6000 and 7500 rpm) can be reproduced using:

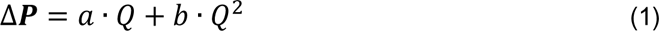

were:

ΔP represents the pressure drop, i.e. the difference between the pressure in the right atrium (RAP)/right ventricle (RVP) and the rotary blood pump pressure (RBPP) for the connection of the inlet cannula and the difference between the pulmonary arterial pressure (PAP) and that of the pump (RBPP) for the outlet cannula. *Q* is the predicted blood pump flow rate (l/min). Table 1 shows the *RBPP* values, as well as *a* and *b* coefficients.

**Table 1.**
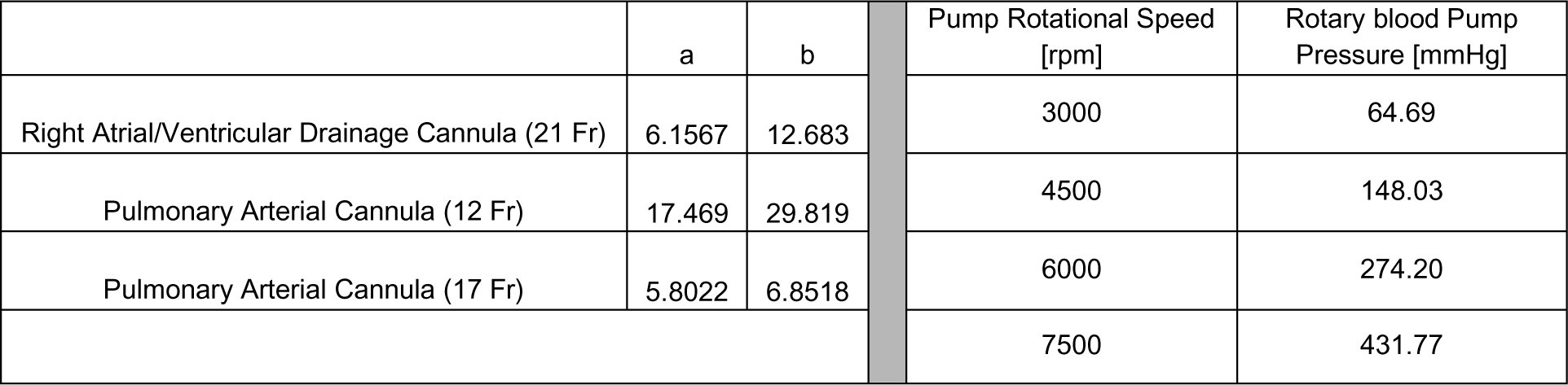

| **Table 1** | **a** | **b** | **Pump Rotational Speed [rpm]** | **Rotary blood Pump Pressure [mmHg]** |
| --- | --- | --- | --- | --- |
| Right Atrial/Ventricular Drainage Cannula (21 Fr) | 6.1567 | 12.683 | 3000 | 64.69 |
| Pulmonary Arterial Cannula (12 Fr) | 17.469 | 29.819 | 4500 | 148.03 |
| Pulmonary Arterial Cannula (17 Fr) | 5.8022 | 6.8518 | 6000 | 274.20 |
|  |  |  | 7500 | 431.77 |

The new ProtekDuo^TM^ module was implemented in CARDIOSIM^©^ using Eqs. 2 and 3 obtained from the experimental “pressure drop-flow rate” waveforms shown in Fig.1. ProtekDuo^TM^ is a dual-lumen cannula: the proximal inflow lumen is positioned in the right atrium; the distal outflow lumen is positioned in the main pulmonary artery. The inflow (29 Fr) and the outflow (16 Fr) port lumens are connected with the TandemHeart^TM^ in the numerical model.

**Figure 1.**
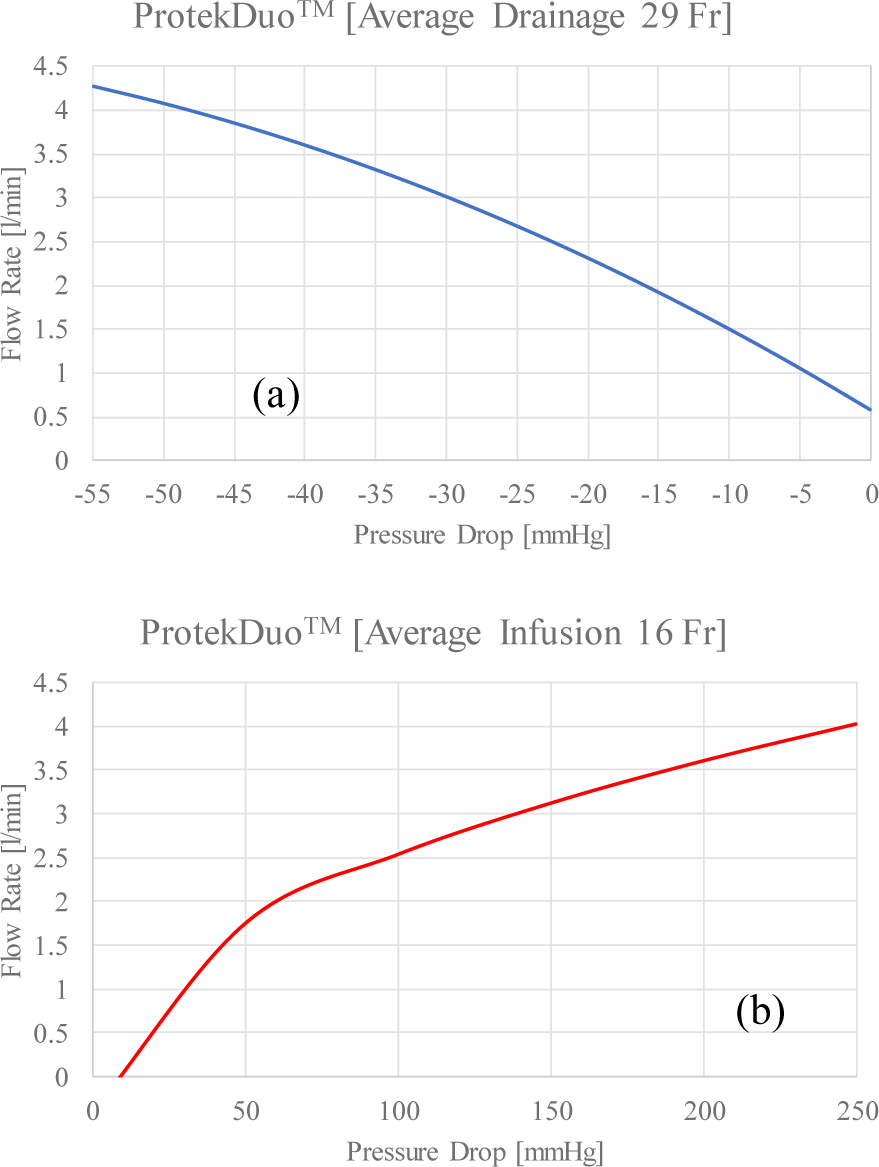
Pressure drop-flow rate waveforms of the proximal inflow lumen positioned in the right atrium (a) and the distal outflow lumen positioned in the main pulmonary artery (b). The waveforms were obtained starting from data published in [27].

The inflow cannula (29 Fr lumen) behaviour is modelled using:

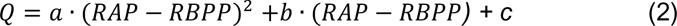

where:

*Q* is the blood flow, *a*=-0.0005557, *b*=-0.09766, *c*=0.5801; *RAP* is the right atrial pressure, *RBPP* is the rotary blood pump pressure, R-square is 0.9999 and adjusted R-square is 0.9997.

The outflow cannula (16 Fr lumen) behaviour is modelled using:

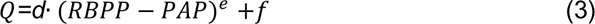

where:

*d*=0.3754, *e*=0.4471 and *f*=– 0.3993, R-square is 1.0 and adjusted R-square is 0.9999.

The coefficients R-square in Eq. 2 and Eq. 3, are all very close to 1 implying a very good fit of the models to the experimental data [28].

For the purposes of our study, the two new modules, which simulate the behavior of the TandemHeart^TM^ pump and the ProtekDuo^TM^ plus TandemHeart^TM^ device, were combined with five modules of the cardiovascular network described in [29-33] as follows:

- Left and right heart including the septum [29,30,33],
- Systemic arterial section [30],
- Systemic venous section [33],
- Coronary circulation [32,33],
- Main and small pulmonary arterial section, pulmonary arteriole, capillary and venous sections [30,33].

Figure 2 shows an electric analogue of the cardiovascular system with the RVAD connected in different configurations.

**Figure 2.**
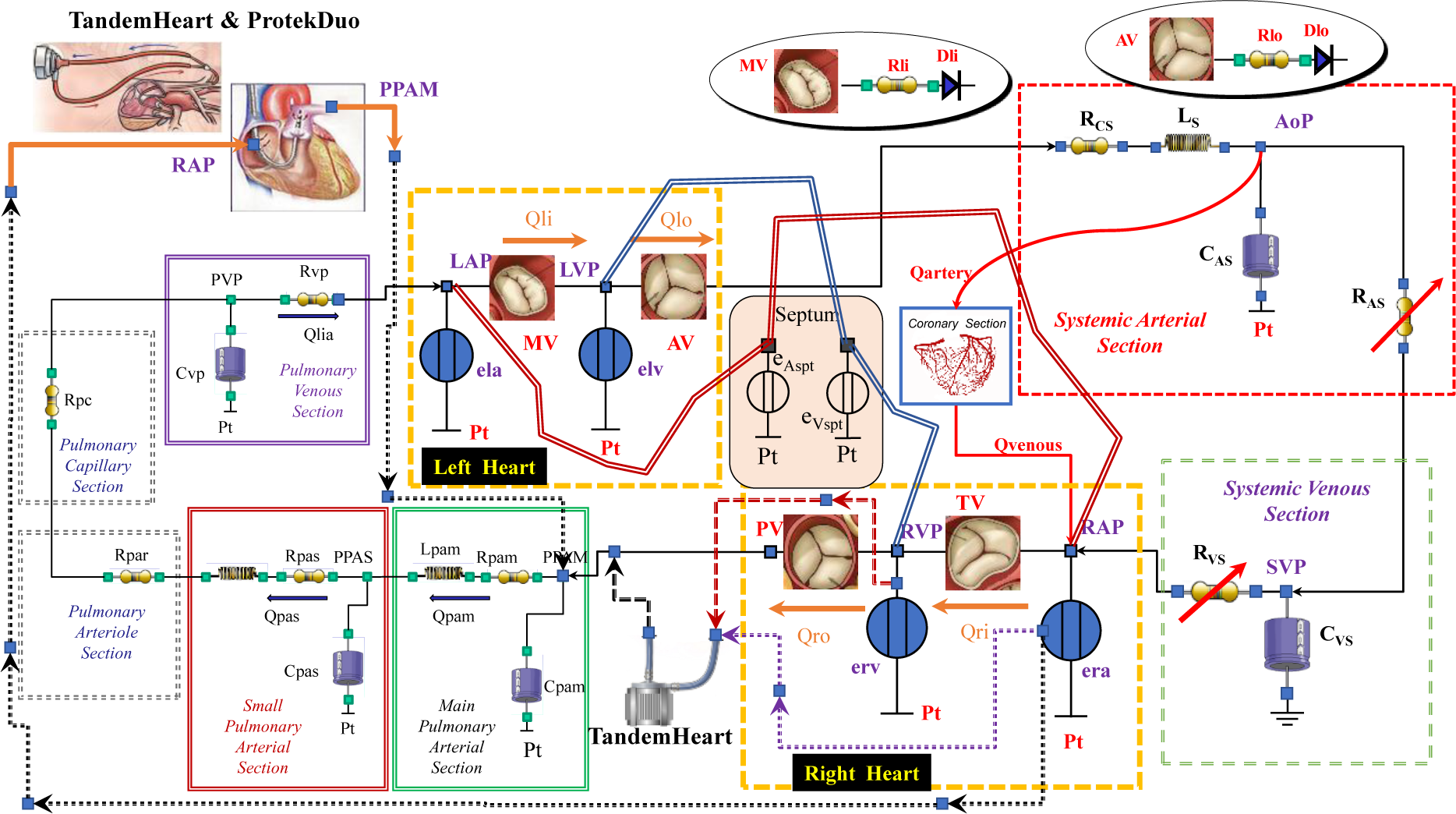
Electrical analogue of the cardiovascular system with the RVAD connected "in series" from the right ventricle to the pulmonary artery and "in parallel" from the right atrium to the pulmonary artery, respectively. All the circuit elements have been described in [29-33]

The heart numerical model, which consists of the left ventricle and atrium (left heart), the right ventricle and atrium (right heart), and the septum, is included in the network configuration assembled for the study. All these elements are described by variable elastance model; elv (erv) represents the left (right) ventricular time-varying elastance; ela (era) is the left (right) atrial time-varying elastance; e_Vspt_ (e_Aspt_) represents the time-varying interventricular (interatrial) septum [29,30].

To simulate acute RV failure, the parameters of CARDIOSIM^©^ were set to achieve a circulatory condition with a mean pulmonary arterial pressure (mPAP) greater than 20 mmHg [34]. Pulmonary arterial hypertension (PAH) with a high mPAP value (> 20 mmHg) was generated by adjusting the mean pulmonary artery and pulmonary arteriole resistances (Fig. 2) [34,35].

Ventricular-arterial coupling (VAC) is a way to determine the optimal power and efficiency of heart performance. The end-systolic elastance (Ees), which is the slope of the end-systolic pressure volume (P-V) relationship [36], and the pulmonary arterial elastance (Ea) [37] are the two parameters that quantify the RV-PA coupling ratio (Ees/Ea). Given that RV-PA coupling measurement is both predictive and diagnostic in PH, the software simulator’s parameters were set to achieve a coupling ratio Ees/Ea=0.56 (Ees/Ea <0.65-0.7 is usually associated with clinical worsening in PAH [38,39]). In this context, the pulmonary capillary wedge pressure (PCWP) was 28 mmHg [12].

Subsequently, we simulated right heart assistance with TH-PD-RVAD and with both TandemHeart^TM^ pump connected "*in series*" (TH-RVADs) and "*in parallel*" (TH-RVADp) mode [40] driven by varying pump rotational speeds; for TH-RVADs and TH-RVADp assistance the arterial (outflow) cannula was set to 12 Fr and 17 Fr, respectively. Under these conditions, we assessed the impact of RVAD support with and without Milrinone administration on the following hemodynamic and energetic variables:

- Mean pulmonary arterial pressure (mPAP),
- Total flow (it’s equal to the RVAD output flow plus the right ventricular output flow),
- Coronary blood flow (CBF)
- Right (left) ventricular end systolic volume RVESV (LVESV),
- Right (left) ventricular end diastolic volume RVEDV (LVEDV),
- Mean right atrial pressure (RAP),
- Pulmonary capillary wedge pressure (PCWP),
- Mean systemic (pulmonary) venous pressure SVP (PVP),
- Mean aortic pressure (AoP), systolic and diastolic aortic pressure,
- Right ventricular-arterial coupling (Ees/Ea),
- Left ventricular-arterial coupling (Ea/Ees),
- Mean left atrial pressure (LAP).

and energetic variables:

- Right (left) ventricular external work EW_R_ (EW_L_),
- Right (left) ventricular pressure volume area PVA_R_ (PVA_L_),
- Right (left) atrial pressure volume area RA_PVA_ (LA_PVA_).

## Results

Figure 3 shows the impact of the TandemHeart^TM^ pump’s assistance on the pressure-volume ventricular loops at 6000 rpm. The RVAD pump was connected "*in parallel*" aspirating blood from the right atrium through a 21 Fr cannula and ejecting it into the pulmonary artery via a 12 Fr cannula (TH-RVADp). The heart rate (HR) was set at 100 beats per minute in both pathological and assisted conditions.

**Figure 3.**
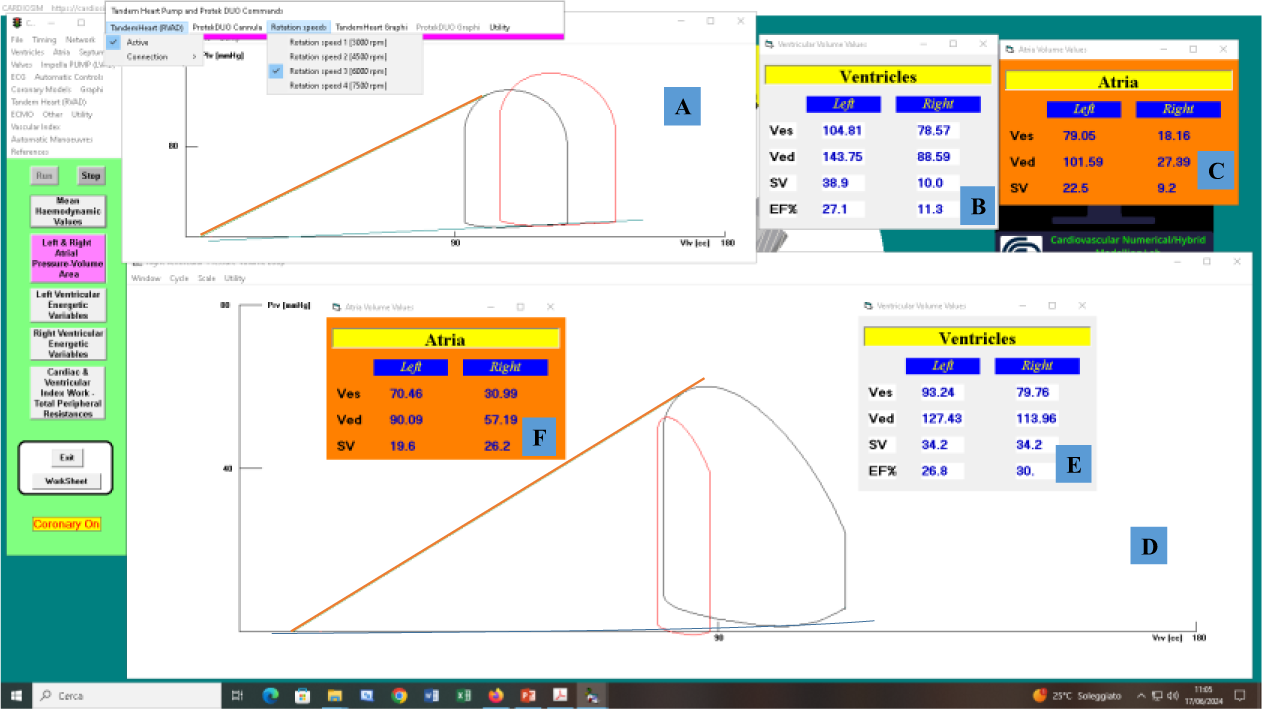
Screenshot generated with CARDIOSIM^©^ software platform. Panel A (D) displays the left (right) ventricular pressure-volume loops simulating RV failure (black line) and the effects of RVAD support (red line). TandemHeart^TM^, driven at 6000 rpm, was used to guarantee RVAD support. The pump takes blood from the right atrium via a 21 Fr cannula and delivers it to the pulmonary artery via a 12 Fr cannula. Panel E (F) displays the pathological left and right ventricular (atrial) end-systolic volume (ESV≡Ves), end-diastolic volume (EDV≡Ved), and stroke volume (SV). Panel B (C) shows the ESV, EDV, and SV recorded when TandemHeart^TM^ was operated "in parallel".

The TH-RVADp reduced the right ventricular end-systolic and end-diastolic volumes (RVESV and RVEDV) (panel D). The right ventricular SV was reduced at 6000 rpm using a right atrial (RA) to pulmonary artery (PA) bypass, according to [4]. The TH-RVAD provided a flow (Q_RVAD_) of about 2.9 l/min, while the right ventricular output flow (Qro) was approximately 1.0 l/min (panel B). The total blood flow (natural blood flow plus RVAD blood flow≅Qlo) was approximately 3.9 l/min. The TH-RVAD decreased both the end-systolic volume and end-diastolic volume for the right atrium (panel C and F). In contrast, the left ventricular stroke volume (SV) with left ventricular end-systolic and end-diastolic volumes (LVESV and LVEDV) increased as a result of the TandemHeart^TM^ support (panel A) [1] according to [1] Panel A shows an increase in left ventricular preload according to [1]. Panels C and F show a decrease in the right atrial pressure volume area (RA_PVA_) on TH-RVADp support.

Figure 4 displays the impact on the left (panel A) and right (panel B) ventricular pressure-volume loops when RVAD support was connected "*in parallel*" and "*in series*" mode with different atrial and arterial cannulae. According to [1] TH-RVADs increased the RVEDV and RVESV (green and blue loops in panel B), whereas RVAD connected in both modes increased LVEDV and LVESV (panel A). RVESV increased and RVEDV decreased when TH-RVADp was connected with a 12 Fr atrial cannula and 17 Fr arterial cannula (purple right ventricular pressure-volume loop in panel B).

**Figure 4.**
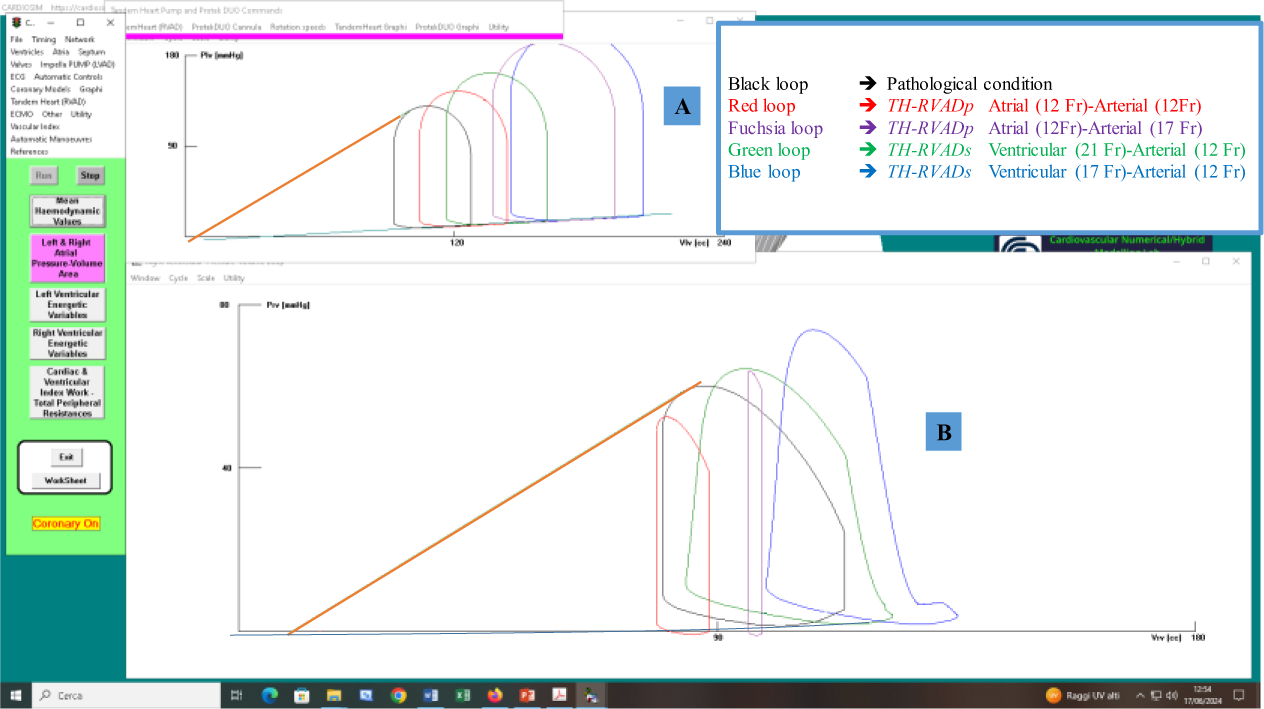
Screenshot generated with CARDIOSIM^©^ software platform. Panel A (B) displays the left (right) ventricular pressure-volume loops simulating RVF (black line) and the effects of RVAD support. Red (purple) loops were generated when the TH-RVADp was connected with atrial cannula of 12 Fr and arterial cannula of 12 Fr (17 Fr). Green (blue) loops were generated when the TH-RVADs was connected with ventricular cannula of 21 Fr (17 Fr) and arterial cannula of 12 Fr. RVAD pump speed was set to 6000 rpm during the whole test.

Figure 5 presents the effects of activating the TandemHeart^TM^ combined with ProtekDuo^TM^ cannula on the pressure-volume ventricular loops at 6000 rpm. The lumen of the ProtekDuo^TM^ inflow cannula (located in the right atrium) is 29 Fr, while the lumen of the outflow cannula (located in the pulmonary artery) is 16 Fr.

**Figure 5.**
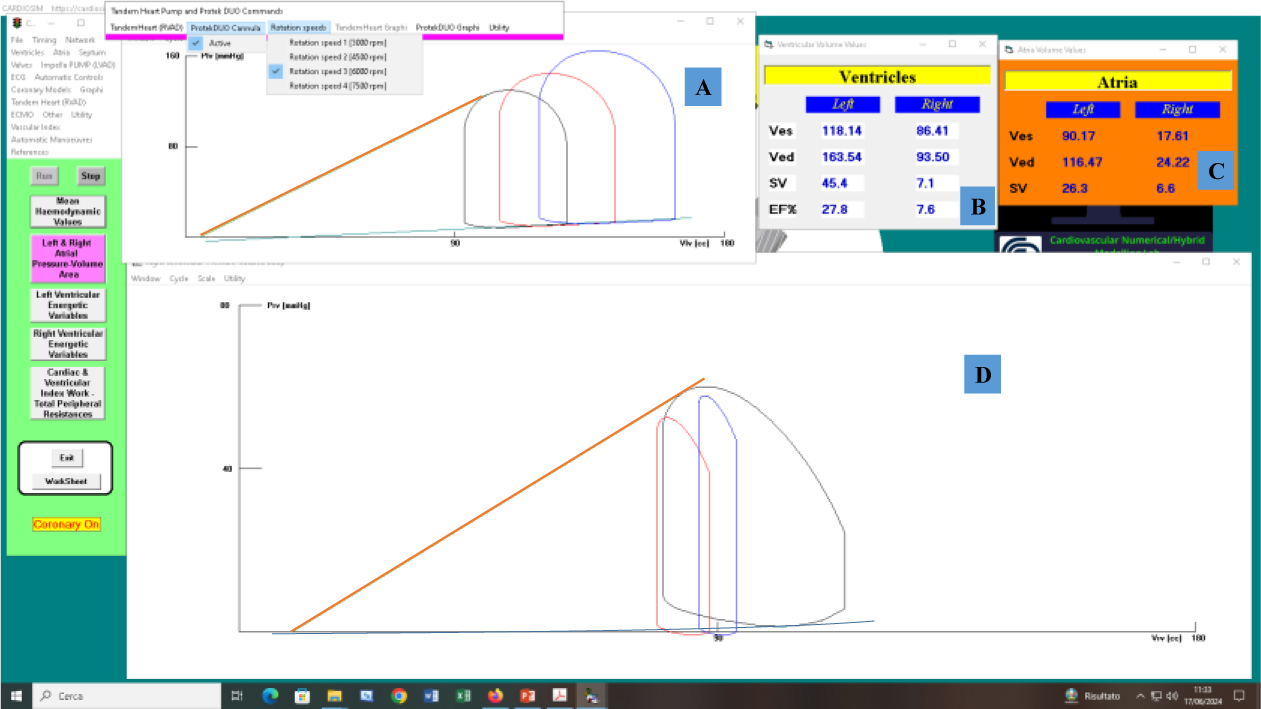
Screenshot produced with CARDIOSIM^©^ software platform. Panel A (D) shows the left (right) ventricular pressure-volume loops simulating RVF (black loops), as well as the effects of TH-RVADp (red loops) and TH-PD-RVAD (blue loops). Panel B (C) presents the RVESV, RVEDV, and SV (for both ventricles and atria) recorded when TH-PD-RVAD was applied.

The TH-PD-RVAD provided a flow of about 3.83 l/min, while the right ventricular output flow (Q_RVAD_) was approximately 0.71 l/min (panel B). The total blood flow (natural blood flow plus RVAD blood flow≅Qlo) was approximately 4.54 l/min. In this scenario, compared to the case described in Fig. 3, the total flow rate had increased by around 16.5%. Comparing panel B, C, E, and F of Fig.3 with panels B and C of Fig.5, it was evident that the use of the ProtekDuo^TM^ cannula led to an increase in the RVESV and a decrease in the RVEDV, resulting in a reduction of the right ventricular external work (EW_R_). Finally, the use of TH-PD-RVAD generated a more significant increase in both LVESV and LVEDV (blue loop in panel A).

Figure 6 shows the relative changes of total blood flow (left side) and the coronary blood flow (right side) calculated in comparison to baseline conditions, when the RVAD was connected “*in parallel*” and “*in series”* mode. TH-RVADp aspirates the blood from the right atrium (with cannula of 21 Fr) and ejects it into the pulmonary artery (with cannula of 12 Fr or 17 Fr).

**Figure 6.**
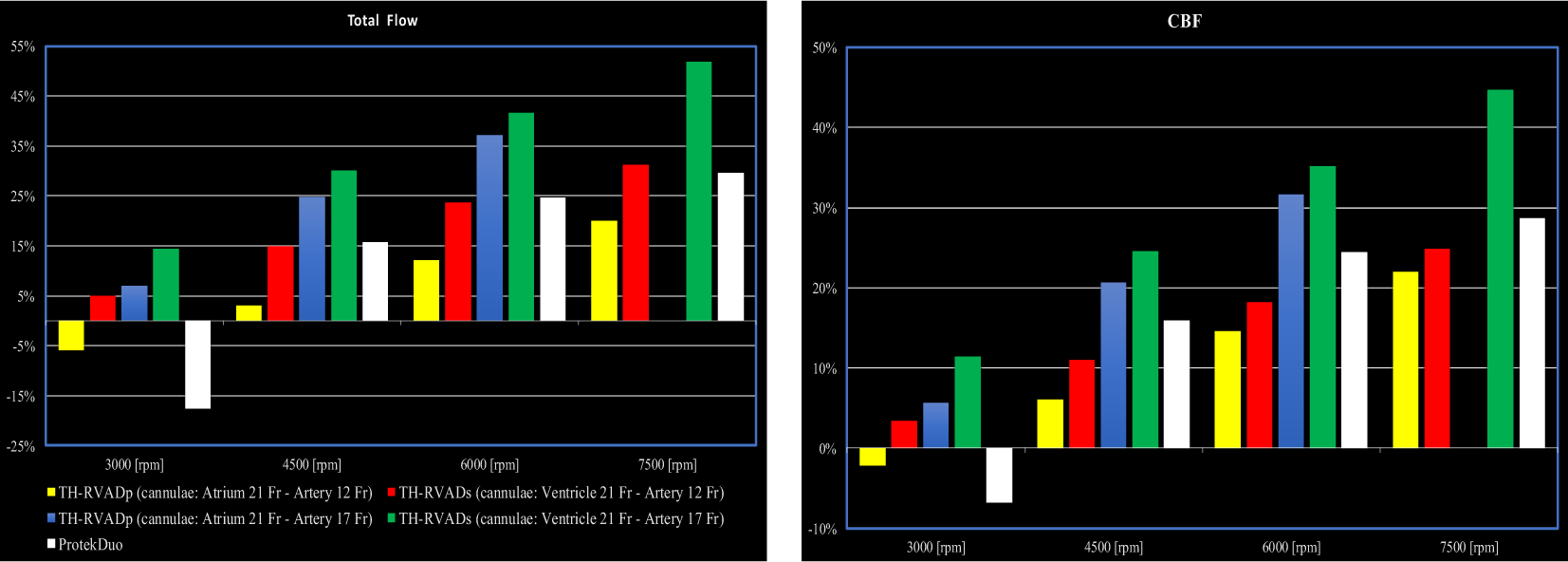
Left (right) panel shows the relative changes of total flow (CBF) calculated in comparison to baseline conditions for TH-RVAD connected “in parallel” (TH-RVADp) and “in series” (TH-RVADs) mode and TandemHeart^TM^ plus ProtekDuo^TM^ cannula. The pump speed was set to 3000, 4500, 6000 and 7500 rpm.

The relative changes of total blood flow and coronary blood flow (CBF) were observed when the TandemHeartT^M^ was connected with ProtekDuo^TM^ cannula (average drainage 29 Fr and average infusion 16 Fr). In the simulated pathological conditions, when the TH-RVADp was connected with a 17 Fr pulmonary artery cannula and activated at 7000 rpm the pump emptied the atrium completely and the software automatically reduced the rotational speed. As a result, no further assessment has been considered for this pump rotational speed. Total blood flow and CBF decreased when the RVAD was connected “*in parallel*" mode with a 21 Fr inflow cannula and a 12 Fr outflow cannula, and when the TH-PD-RVAD support was set with rotational speed of 3000 rpm in agreement to [4]. This is because the inflow cannula empties blood more slowly than the outflow cannula (Fig. 6). The highest relative changes in total blood flow and CBF were recorded when the TH-RVADs was connected with a 21 Fr inflow cannula, a 17 Fr outflow cannula and pump rotational speed at 7500 rpm (green bars) again in agreement to [4].

When the pump speed was set to 4500 rpm, Fig. 7 shows that the TH-RVAD and the TH-PD-RVAD led to percentage increases in RVESV of about 10%. However, larger increases were evident for higher speeds (6000 and 7500 rpm) and connections with 21 Fr inflow and 17 Fr outflow cannula.

**Figure 7.**
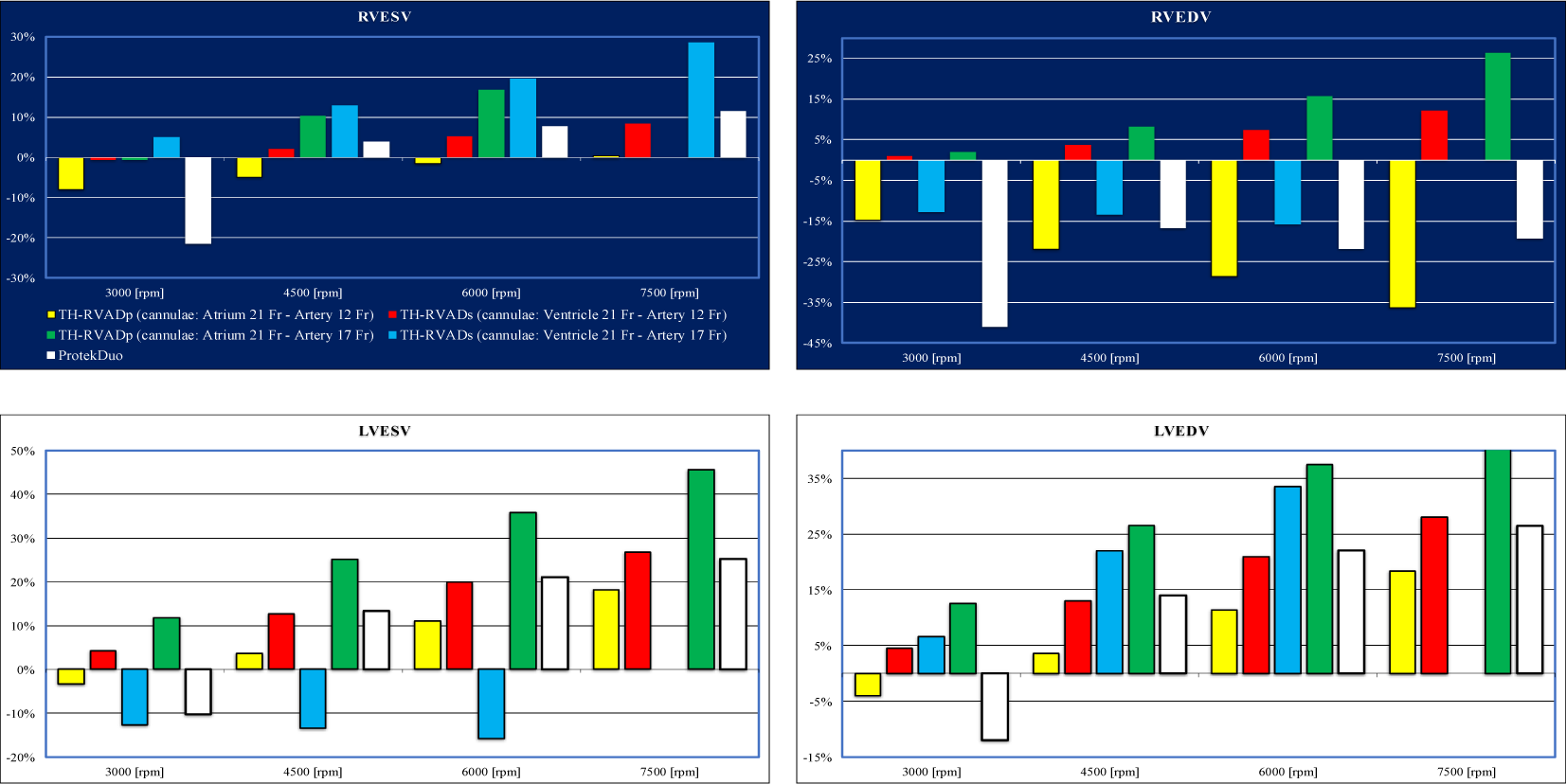
Top left (right) panel shows the relative changes of right ventricular end-systolic (end-diastolic) volume calculated in comparison to baseline conditions for TH-RVADp, TH-RVADs and TandemHeart^TM^ plus dual lumen ProtekDuo^TM^ cannula. Bottom left (right) panel shows the relative changes of left ventricular end-systolic (end-diastolic) volume. The pump speed was set to 3000, 4500, 6000 and 7500 rpm.

According to [41], the top right panel shows how TH-PD-RVAD support reduces the RVEDV.

When TH-RVADp was connected with a 17 Fr outflow cannula, both RVESV and RVEDV increased (green columns in left and top right panels) in agreement to [440].

When TH-RVADs was connected with a 17 Fr outflow cannula, the LVESV decreased, but increased when the pump was connected "*in parallel*" mode (Fig.7 left lower panel). LVEDV increases when the pump speed is equal to or greater than 4500 rpm for TH-RVADp, TH-RVADs and TH-PD-RVAD support (bottom right panel).

The top and bottom left panel of Fig. 8 show the percentage variation of the average pressure in the aorta, pulmonary artery, and left atrium induced by the TH-RVAD connected "*in series*" and "i*n parallel*" with and without ProtekDuo^TM^ dual-lumen cannula positioned in the pulmonary artery.

**Figure 8.**
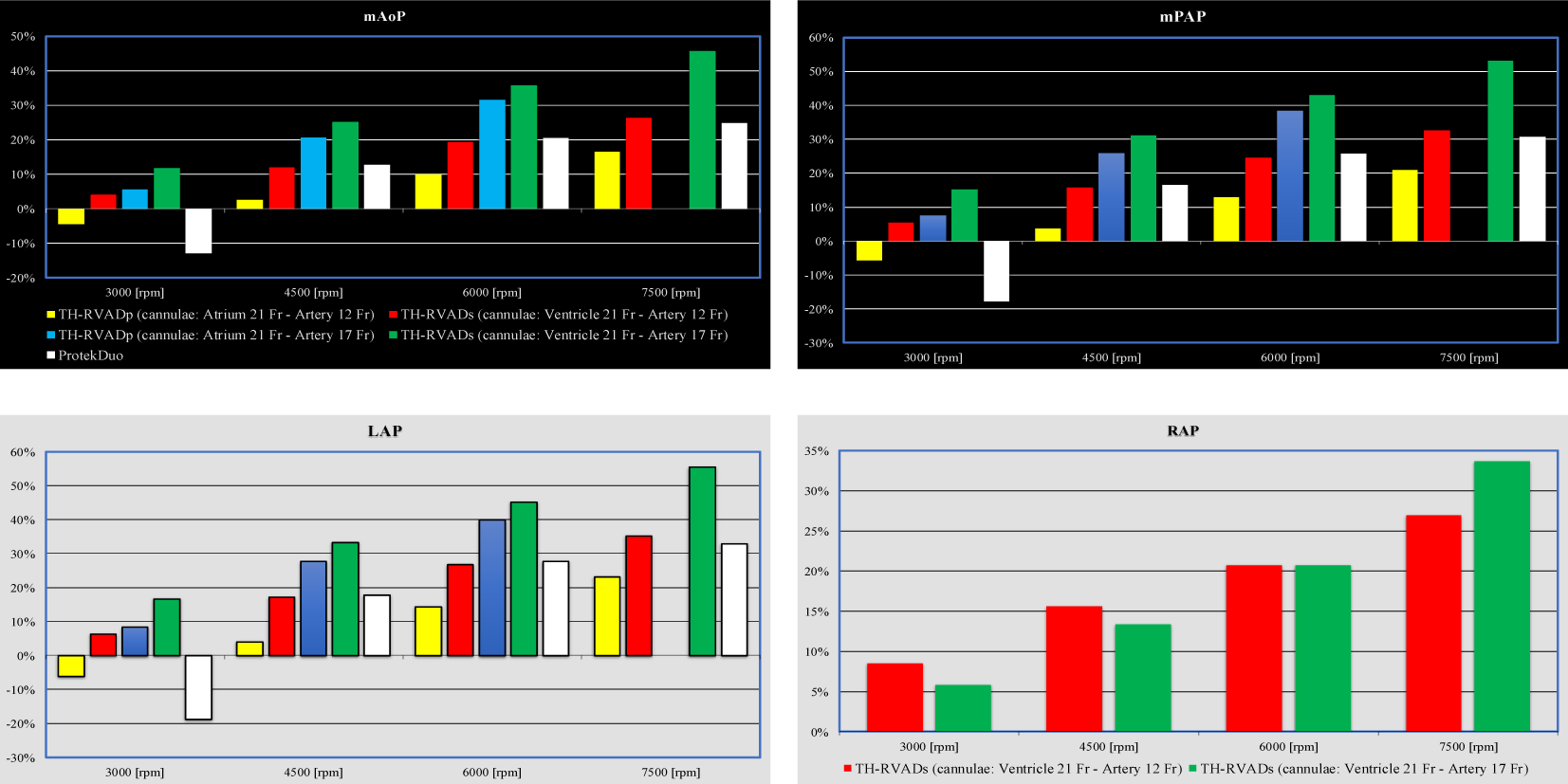
Top left (right) panel shows the relative changes of mean aortic (pulmonary arterial) pressure calculated in comparison to baseline conditions for TH-RVADp, TH-RVADs and TandemHeart^TM^ plus ProtekDuo^TM^ cannula. Bottom left panel shows the relative changes of mean left atrial pressure. Bottom right panel shows the relative changes of mean right atrial pressure for TH-RVADs connected with both 12 and 17 Fr outflow cannula.

When the rotational speed was greater than 3000 rpm, TH-PD-RVAD support increased the mean pulmonary arterial pressure (top right panel), according to [4,41]. TH-RVAD connected in both modes improved mPAP in agreement to [4,40,43].

The bottom right panel (Fig. 8) shows how the percentage variation in mean right atrial pressure increased when the TH-RVADs was connected with 12 and 17 Fr cannulae inserted into the pulmonary artery in agreement to [40]. Finally, TH-RVADp, THRVADs and TH-PD-RVAD increased left ventricular preload (left bottom panel Fig.8) in agreement to [45,46]

Figure 9 shows the effects of RVAD assistance on mean value of PCWP and arteriole pressure (top left and right panel, respectively), as well as systolic and diastolic pulmonary artery pressures (bottom left and right panel).

**Figure 9.**
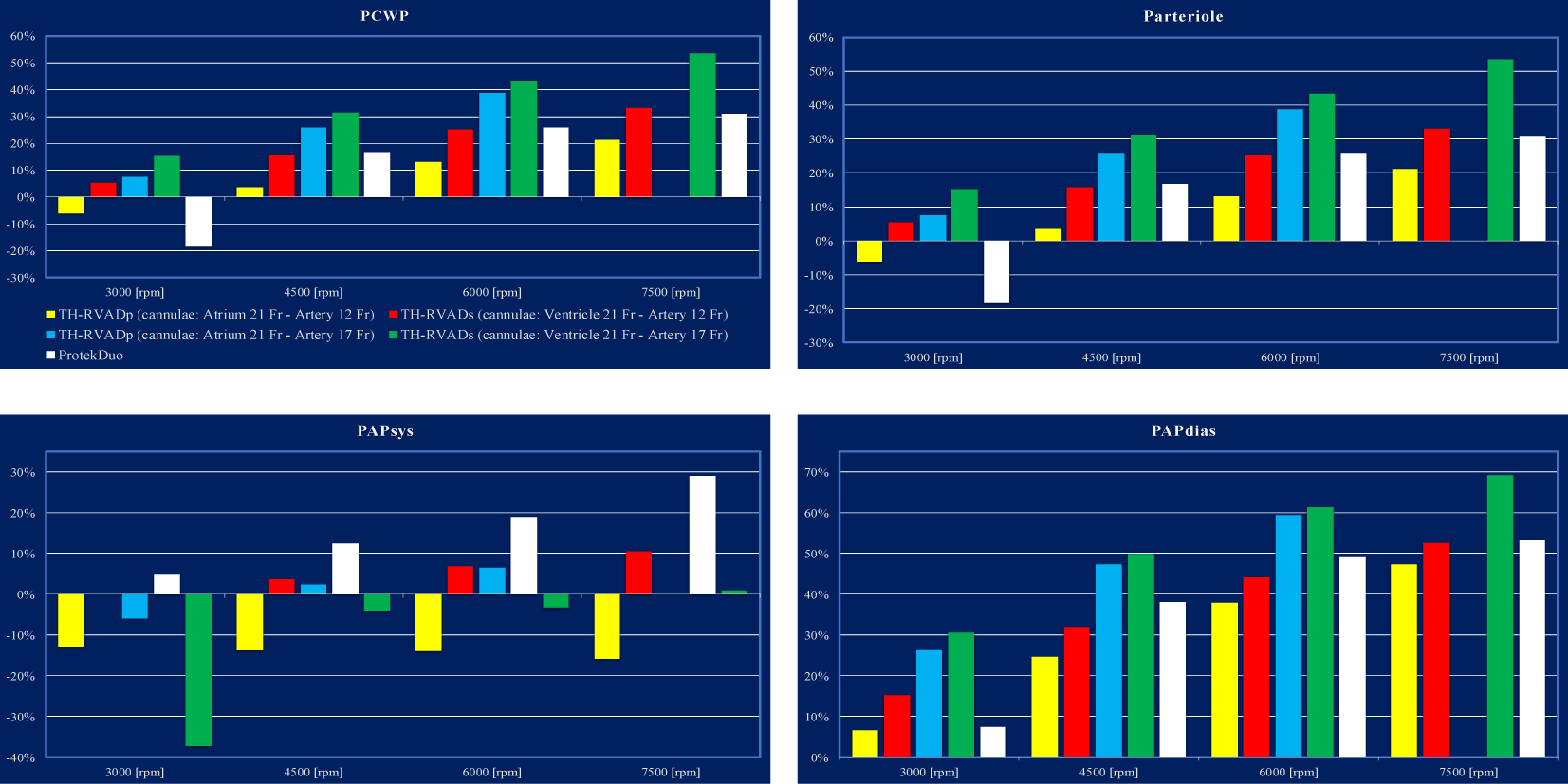
Top left panel shows the relative changes of mean pulmonary capillary wedge pressure (PCWP) calculated in comparison to baseline conditions for TH-RVADp, TH-RVADs and TH-PD-RVAD. Top right panel shows the effects induced on arteriole pressure. Bottom left (right) panel shows the relative changes of systolic (diastolic) pulmonary arterial pressure. The simulations were performed with 12 and 17 Fr pulmonary arterial cannula and the rotational pump speed was set to 3000, 4500, 6000 and 7500 rpm.

When 17 Fr pulmonary arterial cannula was used, the increases in PCWP [42,43], arteriole, and pulmonary diastolic pressures was more evident. When TH-PD-RVAD was applied, the systolic pulmonary arterial pressure increased; however, when TH-RVAD was connected "*in series*" (21 Fr inflow cannula and 17 Fr outflow cannula) and "*in parallel*" (inflow cannula 21 Fr and outflow cannula 12 Fr), the systolic pulmonary arterial pressure decreased [42,43].

Figure 10 presents the effects of the TH-RVADs, TH-RVADp and TH-PD-RVAD on the systemic (top panel) and pulmonary (middle panel) arterial elastance. The bottom panel shows the effects of the TH-RVADs on right (Ees/Ea) and left (Ea/Ees) ventricular-arterial coupling when the pump was connected with 12 and 17 Fr pulmonary arterial cannula

**Figure 10.**
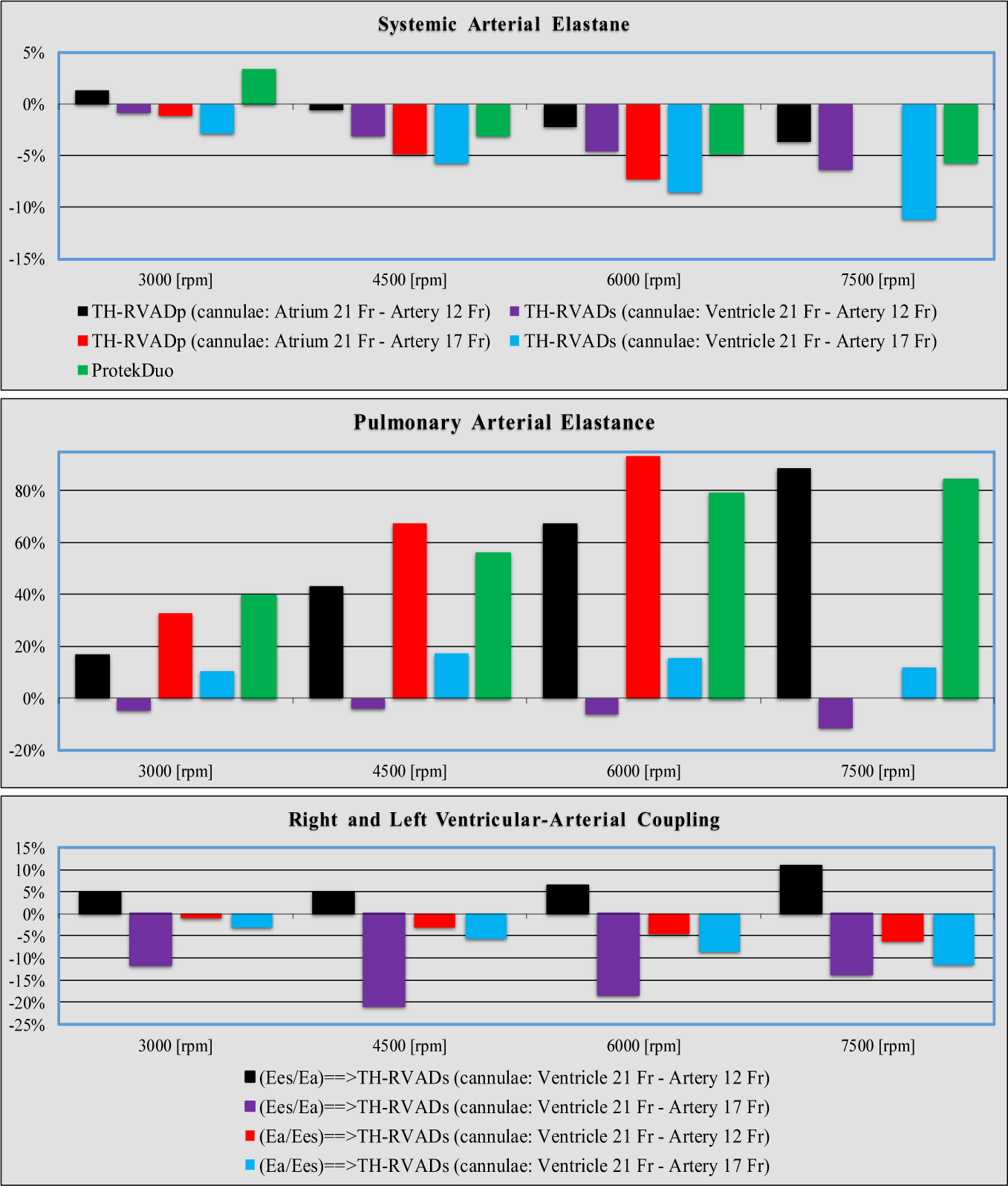
Top (middle) panel shows the relative changes of systemic (pulmonary) arterial elastance calculated in comparison to baseline conditions for TH-RVADp, TH-RVADs and TH-PD-RVAD. Bottom panel shows the effects induced by TH-RVADs on right (Ees/Ea) and left (Ea/Ees) ventricular – arterial coupling. Rotational pump speed was set to 3000, 4500, 6000 and 7500 rpm.

When the pump rotational speed was higher than 3000 rpm, right ventricular assistance reduced systemic arterial elastance (top panel). In contrast, pulmonary arterial elastance (middle panel) increased significantly in all circumstances except for TH-RVADs support connected with a 12 Fr pulmonary arterial cannula. The bottom panel shows the size effect of the pulmonary arterial cannula connected "in series" to the right heart when used with TandemHeart^TM^ support as follows:

- right ventricular-arterial coupling (Ees/Ea) increased in pathological conditions if the pulmonary arterial cannula had a 12 Fr value;
- Ees/Ea decreased in pathological conditions if the pulmonary arterial cannula had a 17 Fr value;
- Left ventricular-arterial coupling (Ea/Ees) decreased in pathological conditions if the pulmonary arterial cannula had both a 12 Fr and a 17 Fr value.

Fig. 11 shows the effects of RVAD support on the energetic variables.

**Figure 11.**
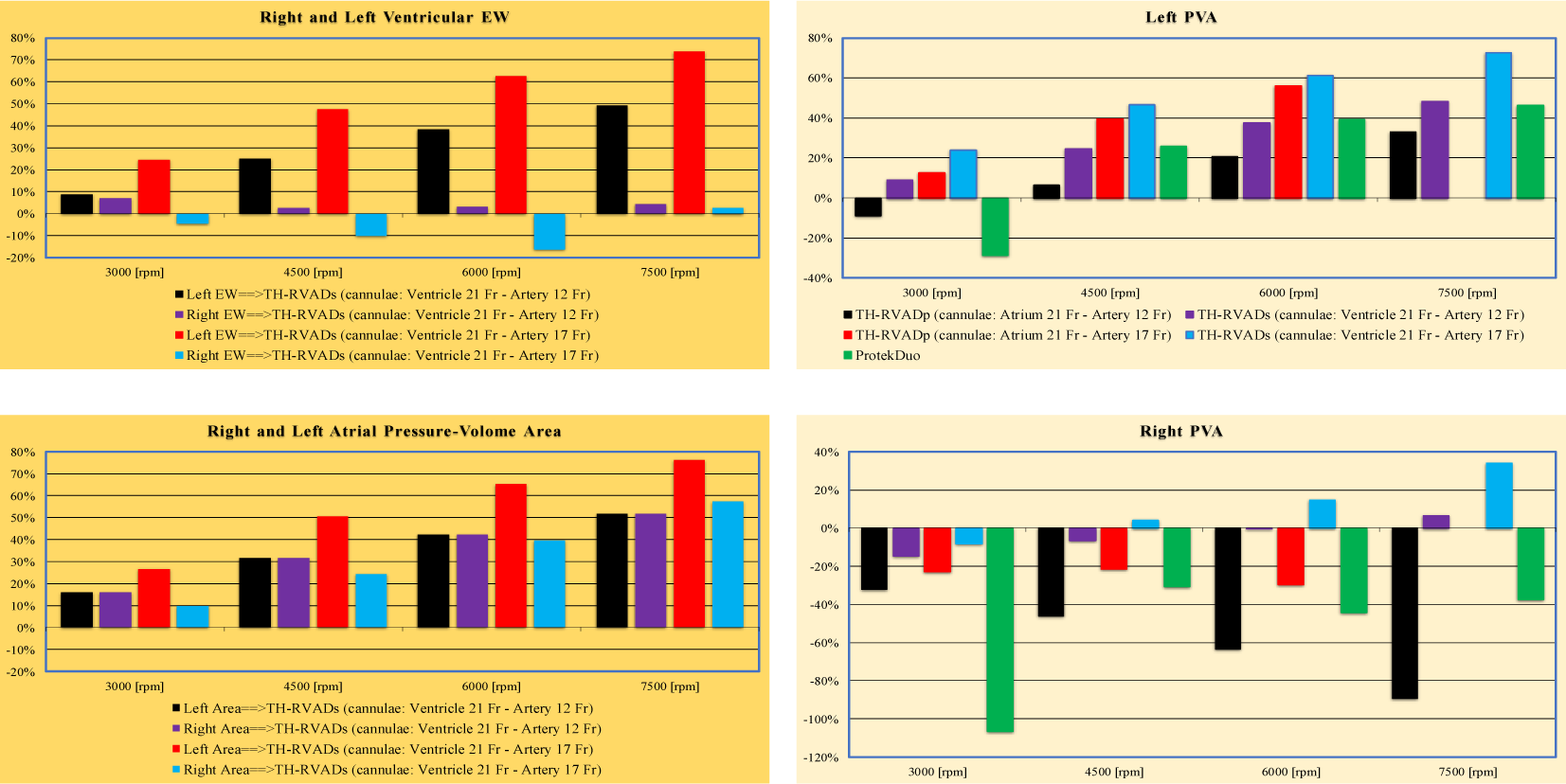
Top left panel shows the relative changes of right (EW_R_) and left (EW_L_) external work calculated in comparison to baseline conditions for TH-RVADs connected with different pulmonary arterial cannulae. Top right (bottom) panel shows the left (right) ventricular pressure-volume area for TH-RVADs,TH-RVADp and TH-PD-RVAD. Left bottom panel shows the right (RA_PVA_) and left (LA_PVA_) atrial pressure-volume area calculated in comparison to baseline conditions for TH-RVADs connected with different pulmonary arterial cannulae.

While a significant rise in EW_L_ was observed in comparison to pathological values, right cardiac support with TH-RVAD connected "*in series*" caused only mild alterations in EW_R_ (top left panel). Regarding the effects of right assistance on PVA_L_, the top right panel illustrates that TH-PD-RVAD and TH-RVADp support with 12 Fr pulmonary arterial cannula reduced PVA_L_ at a rotational speed of 3000 rpm. In any other situation, PVA_L_ increased as a result of both forms of right assistance. Right ventricular PVA (bottom right panel) decreased on TH-PD-RVAD and TH-RVADs support. Only when a TH-RVADp was connected with a 17 Fr pulmonary arterial cannula, an increase in PVA_R_ occur at high pump rotational speed. According to [44,45], PVA is utilized as an indication of oxygen consumption (VO2) in the ventricle. The reduction of PVA_R_ favourably affects oxygen demand and consumption of the right atrium. According to [46] the ideal device would maximally reduce oxygen consumption (demand) by reducing both stroke work and potential energy (and total PVA) while simultaneously augmenting oxygen supply (CBF see right panel of Fig. 6).

Finally, the bottom left panel (Fig.11) shows how RA_PVA_ and LA_PVA_ increased in relation to TH-RVADs rotational speed when the support system was connected with both 12 and 17 Fr pulmonary arterial cannula.

The combined effect on PAP and PCWP (SVP and PVP) induced by assistance with TH-RVADs, TH-RVADp and TR-PD-RVAD and with Milrinone [47,48] is displayed in the top (bottom) panel of Fig. 12. Milrinone is an inodilator that decreases pulmonary resistance and increases cardiac contractility. We estimated a pharmacological dosage in the simulations that would result in a 15% increase in ventricular contractility and a 10% decrease in pulmonary resistance.

**Figure 12.**
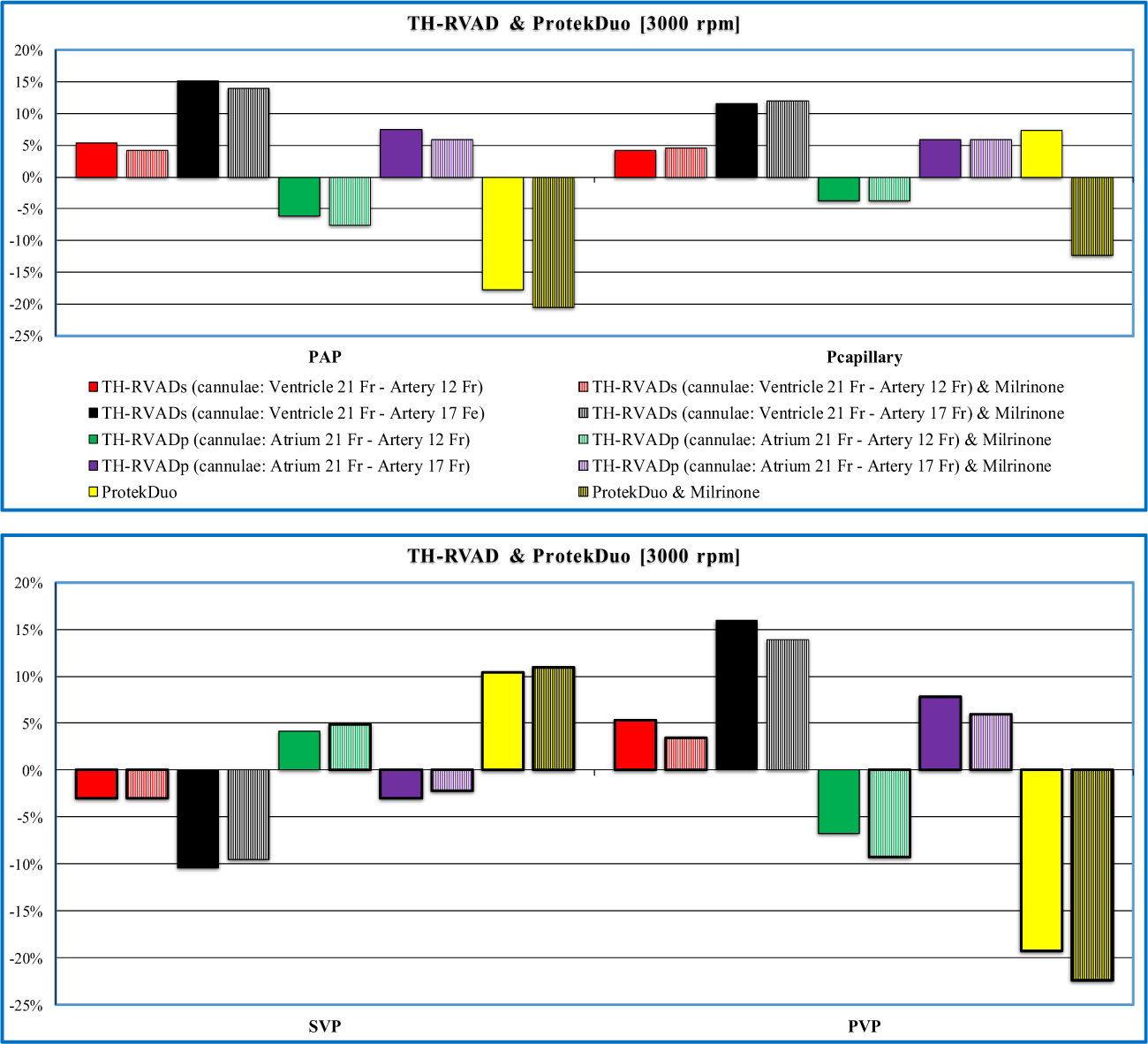
Top panel shows the relative changes of PAP and PCWP (Pcapillary) calculated in comparison to baseline conditions for Milrinone administration during TH-RVADs and TH-RVADp support connected with 12 and 17 Fr pulmonary arterial cannula. Bottom panel shows the systemic venous pressure (SVP) and pulmonary venous pressure (PVP) for Milrinone administration during TH-RVADp (TH-RVADs) with 12 and 17 Fr pulmonary arterial cannula and TH-PD-RVAD support. In all cases, the pump rotational speed was set to 3000 rpm.

In all cases, the pump rotational speed was set at 3000 rpm and the heart rate was maintained constant. According to [49], Milrinone administration reduced PAP while a decrease in PCWP was seen for TH-PD-RVAD support (top panel). Additionally, a reduction induced by Milrinone was observed for PVP [50].

The combined effect on left and right EW following TH-RVADs and TH-RVADp support connected with 12 and 17 Fr pulmonary arterial cannula and with Milrinone administration is displayed in the top panel of Fig. 13. The simulations were performed setting the rotational pump speed to 3000 rpm. Milrinone treatment increased EW_L_ on TH-RVADp and TH-RVADs support whereas an increase in EW_R_ was observed only on TH-RVADp support. The bottom panel of Fig. 13 shows the effects induced by Milrinone administration on PVA_L_ and PVA_R_ following TH-RVADp (TH-RVADs) support with 12 Fr pulmonary arterial cannula and on TH-PD-RVAD assistance. When different RVAD support operated, Milrinone treatment increased PVA_L_ while a slight decrease in PVA_R_ was seen.

**Figure 13.**
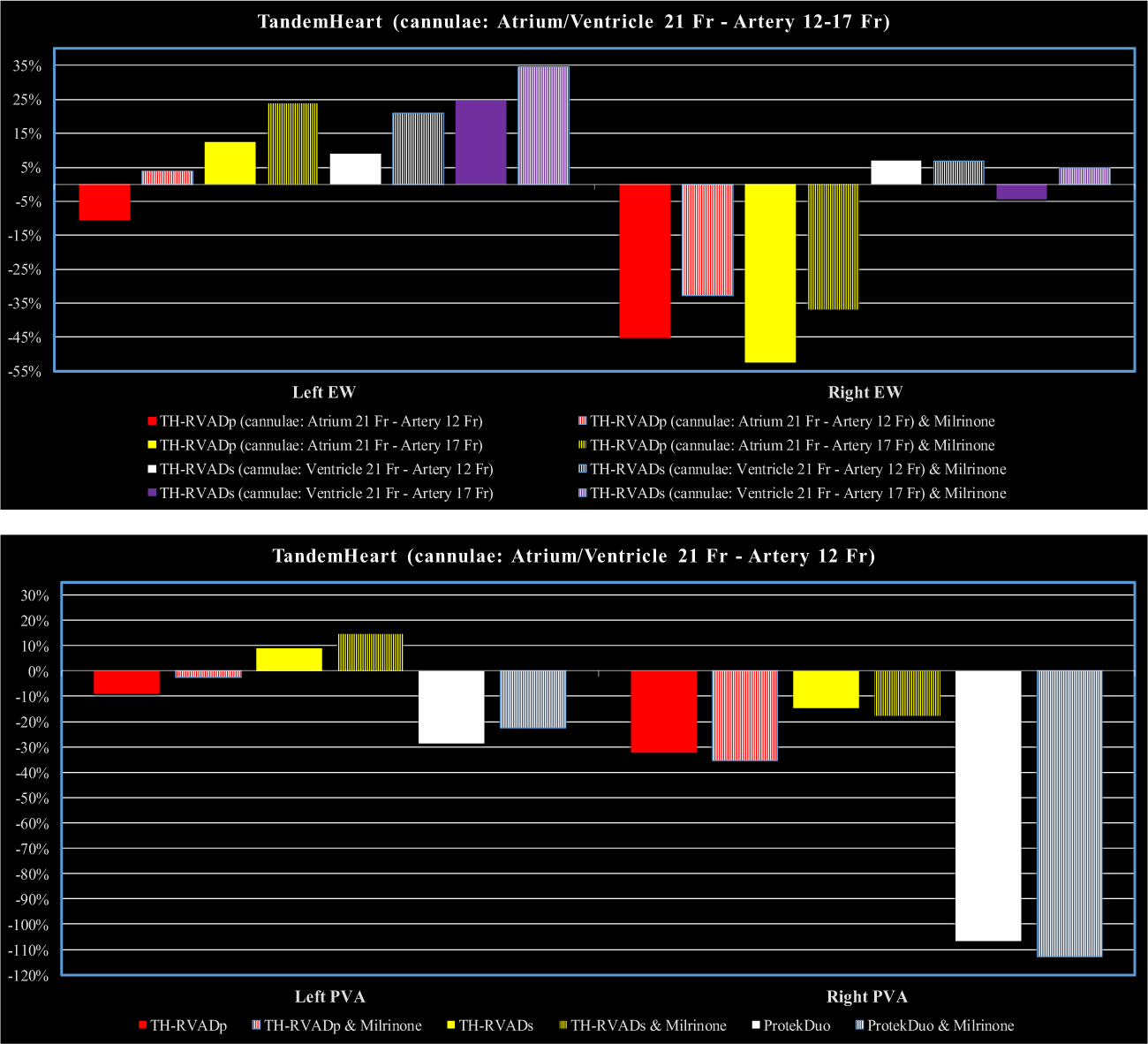
Top panel shows the relative changes of left (EW_L_) and right (EW_R_) external work calculated in comparison to baseline conditions for Milrinone administration during TH-RVADs and TH-RVADp support connected with 12 and 17 Fr pulmonary arterial cannula. Bottom panel shows the left (right) ventricular pressure-volume area for Milrinone administration during TH-RVADp (TH-RVADs) with 12 Fr pulmonary arterial cannula and TH-PD-RVAD assistance. In all cases, the pump rotational speed was set to 3000 rpm.<colcnt=2>

Invasive haemodynamic measurement using a PA catheter is quite helpful for the diagnosis and management of acute RV failure. One way to quantify RV dysfunction is the measurement of the ratio of right atrial (RA) pressure to pulmonary capillary wedge pressure (PCWP). A ratio > 0.86 is highly related to RV failure in the context of acute myocardial infarction [51] whilst a ratio > 0.63 is considered as a predictor of RVF following LVAD insertion [52]. RV stroke work index (RVSWI) is another important haemodynamic parameter to assess RV function [52,53]. Its calculation requires measurement of cardiac output with the Fick method instead of the thermodilution technique, which may underestimate cardiac output in the presence of tricuspid regurgitation [54-57] although its use remains most appropriate in clinical practice [58]. The pulmonary artery pulsatility index (PAPi), defined as [(systolic pulmonary artery pressure – diastolic pulmonary artery pressure)/central venous pressure (right atrial pressure)], is a reliable marker for the prediction of severe right ventricular failure in the setting of acute inferior wall myocardial infarction [59] and following LVAD insertion [53,60]. A PAPi < 1.85 is related to right ventricular failure following LVAD insertion [50] whilst a PAPi < 1 seems related to right ventricular failure in the context of acute myocardial infarction [59]. The presence of pulmonary hypertension may reduce the accuracy of PAPi due to cardiac remodelling [1]. Measurement of the trans-pulmonary gradient, defined as (mean PA – PCWP), can help differentiate right heart failure as a result of left heart failure from diseases of the pulmonary vasculature although the diastolic pressure gradient, defined as mean PAP/PCWP, seems more accurate for the diagnosis of severe pulmonary hypertension [61]. Validated risk-prediction models for right ventricular failure have been proposed. Nevertheless, their overall performance remains poor with limited clinical application [62]. Therefore, a simulation approach in clinical setting with a view to treatment optimisation and outcome prediction may give a different dimension and generate more critical thinking in the context of a multidisciplinary meeting. Lumped-parameter modelling combined with modified time-varying elastance and pressure-volume analysis gives a sufficiently accurate quantitative method applicable within the constraints of the clinical environment. Our results help understand the physiology of the ProtekDuo^TM^ and support the clinical findings [11]. Cannula size, mode of support and pump rotational speed is critical in this setting. The 29-Fr cannula is suitable for patients with a weight less than 45 kg whilst the 31-Fr cannula is suitable for patients with a weight higher than 45 kg. Our simulations focused on the 29-Fr cannula. The parallel mode completely bypasses the right ventricle with favourable effect on preload, ventricular volumes and myocardial oxygen consumption.

## Conclusions

The outcome of our simulations confirms the effective haemodynamic assistance provided by the ProtekDuo^TM^ as observed in the acute clinical setting. A recent systematic review has provided further evidence of its potential for the support of right ventricular failure secondary to different causes [6]. A simulation approach based on pressure-volume analysis combined with modified time-varying elastance and lumped-parameter modelling remains a suitable tool for clinical applications.

## Abbreviation

RVAD (LVAD): Right (Left) Ventricular Assist Device
TH-RVAD: TandemHeart^TM^ in RVAD configuration
TH-PD-RVAD: TandemHeart^TM^ plus ProtekDuo^TM^ in RVAD configuration
TH-RVADs (TH-RVADp): TandemHeart^TM^ connected “*in series*” (“*in parallel*”) mode
PA: Pulmonary artery
LAP (LV): Left atrial pressures (Left ventricle)
RVF: Right ventricular failure
RA-PA (RV-PA): Right atrial (ventricular)-pulmonary arterial
RAP (RVP): Right atrial (ventricular) pressure
PAP (mPAP): Pulmonary arterial pressure (mean pulmonary arterial pressure)
RBPP: Rotary blood pump pressure
PAH: Pulmonary arterial hypertension
VAC: Ventricular-arterial coupling
Ees (Ea): End-systolic (arterial) elastance
PCWP: Pulmonary capillary wedge pressure
SVP (PVP): Mean systemic (pulmonary) venous pressure
CBF: Coronary blood flow
RVESV (LVESV): Right (left) ventricular end systolic volume
RVEDV (LVEDV): Right (left) ventricular end diastolic volume
RAP: Mean right atrial pressure
Aop (mAoP): Aortic pressure (mean aortic pressure)
Ees/Ea (Ea/Ees): Right (left) ventricular-arterial coupling
EW_R_ (EW_L_): Right (left) ventricular external work
PVA_R_ (PVA_L_): Right (left) ventricular pressure volume area
RA_PVA_ (LA_PVA_): Right (left) atrial pressure volume area
EDV≡Ved (ESV≡Ves),: End-diastolic (systolic) volume
SV: Stroke volume
QRVAD: TH-RVAD pump flow
Qro (Qlo): Right (left) ventricular output flow
PAPsys (PAPdias): Systolic (diastolic) pulmonary arterial pressure
PVA (EW): Pressure volume area (external work)
VO2: Oxygen consumption
RVSWI: Right ventricular stroke work index
PAPi: Pulmonary artery pulsatility index

